# Proposal of a simple open-source quantitative tribometric assay and its implementation for the assessment of the effects of redox-related alterations on the lubrication capacity of a commercial water-based lubricant gel

**DOI:** 10.1101/2022.07.27.501731

**Authors:** Jan Homolak

## Abstract

**Background:** Understanding molecular and biochemical mechanisms affecting biotribological properties of tissues, biological fluids, and drugs may accelerate the invention of novel drug formulations and targets and facilitate the discovery of etiopathogenetic mechanisms. Unfortunately, biotribometric equipment is expensive and unaffordable. The first aim was to assess whether PASTA, an open-source platform based on a hacked kitchen scale, can be adapted for the analysis of biotribometric properties. The second aim was to demonstrate the use of PASTA by studying the effects of oxidation on the lubricating properties of the commercial water-based lubricant.

**Methods:** The PASTA platform was adapted using a custom-made tribometric adapter attached to the bottom of the hacked kitchen scale connected to a computer via the NodeMCU ESP-32S. A commercial water-based lubricant was oxidized with 2,2’-azobis(2-amidinopropane) dihydrochloride and/or protected against oxidation with glutathione. The samples were analyzed using a simple tribometric assay with the PASTA platform and the ORP-146S redox microsensor.

**Results:** A mastPASTA custom-made adapter enables the use of PASTA for reliable quantitative tribometric studies. Oxidation of the commercial water-based lubricant does not reduce its lubrication capacity, however, the addition of the antioxidant glutathione protects against the loss of its lubrication capacity upon exposure to air by a redox-unrelated mechanism.

**Conclusion:** PASTA can easily be adapted for studying tribological properties. The effects of the addition of antioxidants to water-based vaginal lubricants should be explored as a possible way to enhance durability and efficacy and increase their health benefits.

## Introduction

Biotribology is an interdisciplinary scientific field focused on studying the properties of interacting surfaces in relative motion inside a complex biological system. The principles of biotribology can be applied to many different biomedical scenarios – e.g. studying the wear, tear, and regenerative properties of synovial joints[1], understanding the physiological and pathophysiological properties of mucus[2–4] or developing novel drug delivery systems[5]. A better insight into the molecular and biochemical mechanisms responsible for the modulation of biotribological properties of tissues, biological fluids, and drugs may accelerate the invention of new drug targets[6] and more effective drug formulations[7], and facilitate the discovery of novel etiopathogenetic mechanisms[4,8–10]. Unfortunately, studying tribometric properties of tissues, biological fluids, and drugs usually requires sophisticated and expensive laboratory equipment unaffordable to many.

Recently, an increase in the availability of low-cost electronics and the growing accessibility of educational material and literature on the topic encouraged many scientists to join the open-science community and work together on the development of open-source scientific instruments[11–17]. The open-source tools offer many advantages in comparison with the commercial instruments: i) they can usually be made with easily available equipment and on the cheap; ii) they are characterized by a flexible and modular design at the level of both software and hardware which makes them easily adaptable to a wide variety of specific use-case scenarios; iii) they are supported by a growing community of researchers passionate about what they are doing.

The first aim of the present study was to assess whether PASTA – an open-source platform based on a hacked kitchen scale interfaced with a computer originally introduced by Virag et al. [14] can be modified for the assessment of tribological properties. The second aim was to demonstrate the use of PASTA with a simple biotribometric study by assessing whether oxidation affects the lubricating properties of a commercial water-based lubricant gel.

Oxidation plays a key role in the aging and deterioration of the lubricating capacity of many lubricants (e.g. [18]), and antioxidants are considered to be important additives in water-based lubricants [19]. Nevertheless, the importance of redox alterations has to the best of my knowledge never been explored in the context of the lubricating capacity of water-based lubricants for personal use. Consequently, a working hypothesis was postulated that the addition of antioxidants may increase the resilience of water-based lubricants for personal use to environmental conditions that take a toll on their lubricating properties (e.g. exposure to air). In case the working hypothesis proves to be correct it has many real-life implications as antioxidants may also provide some health benefits in women suffering from vaginal dryness (VD) - a medical condition for which they are commonly used as a symptomatic treatment. VD is a prevalent disorder characterized by insufficient lubrication of the vagina[20] that affects the quality of life and poses an economic burden in both premenopausal and postmenopausal women [21]. In postmenopausal women, VD seems to be a direct consequence of the reduced levels of estrogen associated with depletion of antioxidants and oxidative stress[22–24] and vaginal administration of royal jelly with antioxidant properties has shown some benefits regarding quality of life and vulvovaginal atrophy [25] suggesting that local administration of antioxidants should be examined as a treatment strategy. The potential benefits of locally administered antioxidants may be especially relevant during sexual intercourse (e.g. the period when lubricants are standardly used) as increased shear stress has been associated with increased cellular oxidative stress[26,27].

## Materials and methods

### Sample preparation

Durex Play Saucy Strawberry Lube (Reckitt Benckiser, UK) was used as a standard sample for the proof of concept study. Based on the list of ingredients the lube contains glycerin, H_2_O, propylene glycol, hydroxyethylcellulose, benzoic acid, sodium saccharin, sodium hydroxide, and an additional undisclosed, aroma” ingredient. To test the effects of air exposure, oxidation, and antioxidant protection the lube (500 μl) was mixed with either i) 200 μl of ddH_2_O (CTR); ii) 100 μl of the 92 mM solution of 2,2’-azobis(2-amidinopropane) dihydrochloride (AAPH) + 100 μl of ddH_2_O (AAPH); iii) 100 μl of the 100 mM solution of the reduced glutathione + 100 μl of ddH_2_O (GSH); iv) 100 μl of AAPH + 100 μl of GSH. All samples were mixed and left at room temperature for 120 min. After 120 min, an additional sample was prepared for tribological analysis by mixing 500 μl of the lube with 200 μl of ddH_2_O (CTR_fresh).

### The oxidation-reduction potential and the pH

The oxidation-reduction potential (ORP) was measured to ensure the samples were oxidized in the presence of AAPH and protected against oxidation in the presence of GSH. The analysis was conducted using the redox microsensor system ORP-146S (Lazar Research Laboratories, Los Angeles, USA) with a platinum sensing element, Ag/AgCl reference, and KCl used as a filling solution as described previously[28,29]. Both the ORP and the pH values were obtained with the 6230 N Microprocessor meter (Jenco Instruments, San Diego, USA).

### Evaporation

Evaporation was assessed using the gravimetric method. The samples were weighed before and after the incubation, and the mass of evaporated fluid was calculated by subtracting the post-incubation weight from the pre-incubation weight measured using the analytical balance (Mettler Toledo, Columbus, Ohio, SAD).

### A tribometric adaper for PASTA

The platform for acoustic startle (PASTA) is an open-source device originally developed for measuring acoustic startle and prepulse inhibition sensorimotor gating by Virag et al[14] and later repurposed for quantification of grip strength[15]. The platform consists of a hacked kitchen scale connected to a readily-available affordable controller circuit board relying on the modified HX711 integrated circuit interfaced with a microcontroller (NodeMCU ESP-32S board) that acts as a communication bridge and enables real-time data processing, visualization, and long-term storage on the computer. A general-purpose PASTA script (PASTAgp) [30] was used for data acquisition. For detailed instructions on how to make PASTA please see [14,15,31].

A custom-made tribometric adapter, *Multifunctional Adapter for Screening Tribometry* for PASTA-mastPASTA was designed to enable using PASTA load cells for the acquisition of time series data of the vertical pulling force applied to the tubing system attached to the bottom of the standard PASTA platform placed on a hollow bench. Briefly, a hollow cylinder was fixed onto the bottom of the PASTA platform using two parallel polystyrene frames connected with nuts and bolts. Two small diametrical holes were drilled in the hollow cylinder to enable attachment of the 4 cm long tubing with an inner diameter of 4 mm with a pin on the inside of the cylinder. The polyvinyl chloride (PVC) tubing was marked with a permanent marker and the inner surface of the tubing was used as the interface for the tribometric analyses following the insertion of the narrow PVC tubing with an outer diameter of exactly 4 mm.

### Analysis of the association between surface area and peak resistance

The association between the contact area of the convex and concave PVC surface and the maximal mastPASTA-recorded resistance upon application of a dynamic (vertical) force at a 90° contact angle was measured using a pair of 4 cm long 4 mm inner diameter and 4 cm long 4 mm outer diameter PVC tubes with 50 μl of the normal lube administered onto the concave surface with a trimmed tip pipette. After the lube was administered the inner tubing was inserted 10 times in a way that the maximal achieved surface contact area was ~ 377 mm^2^. After the lubricant gel was spread evenly, the outer tube was connected to the mastPASTA adapter with a pin. The inner tubing was inserted until the marking representing the contact area of 125.66, 251.33, or 376.99 mm^2^ was reached. The inner tubing was pulled vertically (at a 90° contact angle) and PASTA recorded time-series force profiles. The procedure was repeated 6 times for each contact surface area to assess linearity. The peak resistance force for each trial was determined from PASTA force profile plots in R using the plotly package[32].

### Tribometric analysis of the commercial water-based lubricant gel

Tribometric analysis was conducted by i) measuring peak resistance achieved by the contact of the pre-treated convex and concave PVC surface upon application of a dynamic (vertical) force at a 90° contact angle using mastPASTA; ii) by measuring capillary movement using 100 mm long melting point glass capillary tubes with a 0.5 mm diameter; iii) by measuring the movement of the gel on a clean borosilicate surface at a 90° angle.

For the analysis of the peak resistance using the mastPASTA platform, pairs of 4 cm long 4 mm inner diameter and 4 cm long 4 mm outer diameter PVC tubes were prepared for each sample. 20 μl of each sample (CTR, AAPH, GSH, AAPH+GSH) was deposited inside the outer tubing and spread evenly by inserting the inner tubing until the 2 cm marking corresponding to the 251.33 mm^2^ contact surface. The outer tubing was connected to the mastPASTA adapter, the inner tubing was inserted until the 2 cm marking and pulled downwards. The whole procedure was repeated 10-16 times for each sample.

The capillary movement was estimated by placing a 100 mm long melting point glass capillary tube with a 0.5 mm diameter in the sample vertically. The tube was gently adjusted every 60 seconds to avoid biased placement inside the solution. After 300 s the tube was carefully removed from the solution. The same procedure for repeated 3 times for each sample, the tubes were photographed together, and standardized column height for each tube was calculated using Fiji.

The movement of the samples on a clean borosilicate surface placed at a 90° angle was estimated by placing 10 μl of each sample on the microscope slide. The slide was fixed at a 90° angle, and the movement was recorded using Samsung S20 FE (Samsung, Suvon, South Korea). Four samples (CTR, AAPH, GSH, AAPH+GSH) were analyzed in each trial, and the whole procedure was repeated 5 times. The videos were analyzed in Fiji by calculating the total vertical distance covered (maximal length with respect to the movement vector), and distance covered by the sample front at t=0, 5, 10, 15, and 20 s.

### Data analysis

Data were analyzed in R (4.1.3). The tribometric data was reported as population means with confidence limits obtained by non-parametric bootstrapping without assuming normality. The mastPASTA was first transformed from .pasta to .csv and peak values were extracted using the plotly package. Log-transformed peak PASTA values were used as the dependent variable and treatments (CTR, AAPH, GSH, AAPH+GSH) as the independent variable. Model assumptions were checked using visual inspection of the residual and fitted value plots and model outputs were reported as ratios of estimates marginal mean point estimates with corresponding 95% confidence intervals. The Tukey method was used for adjusting the comparisons of a family of 5 estimates. For the tribological analysis of the movement of the gel on a clean borosilicate surface at a 90° angle, the distance moved was used as the dependent variable, the treatment, sampling time-point, and their interaction were modeled as independent fixed effects variables, and the sample was modeled as the independent random effect in the linear mixed model. Assumptions were checked by visual inspection of the residual and fitted value plots, and point estimates were reported with their respective 95% confidence intervals.

## Results

### mastPASTA can be used to quantify lubricating properties

A proof-of-concept experiment presented here demonstrates that PASTA can be easily modified into a platform suitable for the quantitative assessment of tribological properties. Here, a simple adapter (mastPASTA) is proposed in order to enable using the kitchen scale load cells to record forces made by friction between the concave and the convex surface of two complementary PCV tubes fixed on the bottom of the kitchen scale placed on top of a hollow table (Fig 1A-C). The proposed platform can be used to quantitatively assess lubricating properties based on the expected reduction in friction between the PVC tubes the following administration on the contact surface area. The pilot experiment with the *Durex Play Saucy Strawberry Lube* deposited on the concave surface of the PVC tubing demonstrates that the proposed system can be used to test lubricating efficacy by demonstrating the linear association between the maximal recorded resistance and the contact surface area of the tubing (Fig 1D).

**Fig 1.**
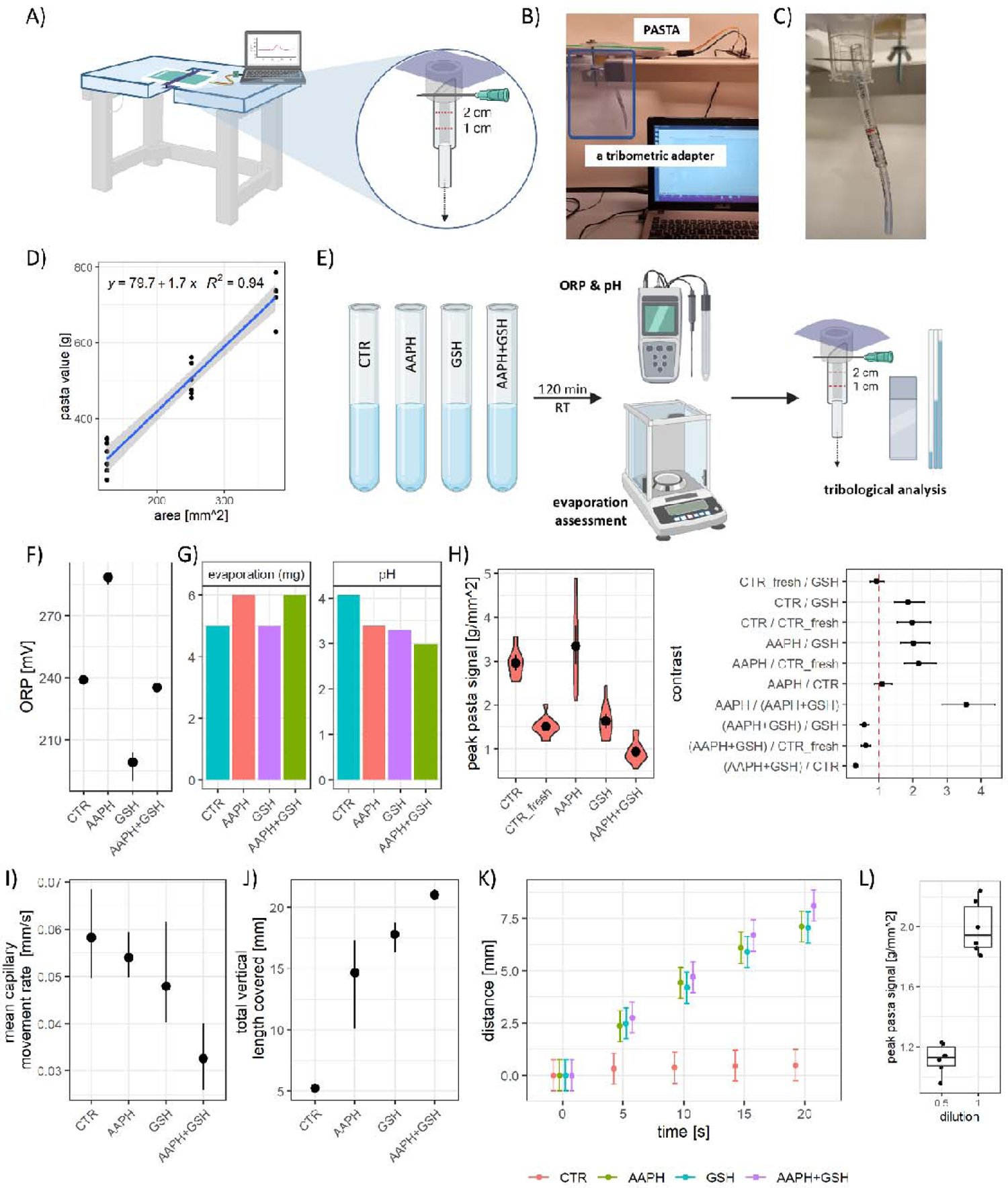
A schematic presentation of the *Multifunctional Adapter for Screening Tribometry* for PASTA (mastPASTA) and the results from the study on the effects of redox-related alterations on the lubrication capacity of the commercial water-based lubricant gel. A) A schematic representation of the PASTA platform with the mastPASTA adapter set up for the quantitative tribological assessment. B) A photograph of the PASTA setup with a mastPASTA tribometric adapter. C) A close-up of the mastPASTA tribometric adapter. D) The association between the absolute contact surface area and the PASTA-derived maximal resistance upon application of a dynamic (vertical) force at a 90° contact angle. E) An overview of the experimental design for the study on the effects of redox-related alterations on the lubrication capacity of the commercial water-based lubricant gel. F) The oxidation-reduction potential (ORP) of samples from E confirms the assumed altered redox status of the lubricant gel. G) Evaporation of fluid during the incubation, and pH value of samples with altered redox status. H) Violin plots of the PASTA-derived maximal resistance upon application of a dynamic (vertical) force at a 90° contact angle upon lubrication of the concave surface with the control and treated samples (left) and point estimates of ratios of least square means accompanied by corresponding 95% confidence intervals illustrating differences in lubricating properties introduced by alteration of the redox status of the water-based commercial lubricant gel. I) Point estimates and non-parametric bootstrapping-derived confidence limits of mean capillary movement rate of the samples. J) Point estimates and non-parametric bootstrapping-derived confidence limits of mean total vertical distance covered on the borosilicate surface at a 90° angle. K) The temporal pattern of the movement of samples on the clean borosilicate surface fixed at a 90° angle. L) CTR – control; AAPH - 2,2’-azobis(2-amidinopropane) dihydrochloride-treated sample; GSH – glutathione-treated sample; AAPH+GSH – sample treated with both AAPH and GSH.

### The addition of exogenous GSH to the lubricant prevents the decrement in its lubrication efficacy upon exposure to air, but likely not due to its antioxidant properties

To demonstrate its usefulness in conducting simple tribometric analyses following the installment of the mastPASTA adapter PASTA was utilized to study the effects of redox-related alterations on the lubrication efficacy of the *Durex Play Saucy Strawberry Lube* (Fig 1E). Aliquots of the lube were incubated for 120 minutes with i) vehicle (as a control); ii) AAPH – to induce oxidation; iii) AAPH + GSH – to try to counteract the oxidation by AAPH; iv) GSH – to control for the redox-unrelated effects of GSH. The ORP analysis confirmed that the incubation with AAPH and GSH induced a discernible pro- and anti-oxidative shift respectively (Fig 1F). Furthermore, the lubricant incubated simultaneously with both AAPH and GSH showed a minimal alteration of the ORP indicating that oxidation by AAPH was successfully counteracted by the addition of GSH (Fig 1F). Since the samples were exposed to air during the incubation, the evaporation was analyzed to prevent potential bias that may have been introduced by the unforeseen effects of exogenous substances on the lubricant desiccation rate. The gravimetric analysis confirmed that there was no pronounced effect of the added substances on the lubricant desiccation rate during the incubation (Fig 1G). Additionally, the pH of the samples was determined to take into account potential differences in tribological properties that can be attributed to pH rather than redox alterations introduced by the treatments (Fig 1G). Tribological analysis of the samples with the mastPASTA showed that AAPH-mediated oxidation had a small and inconsistent effect on the lubricating efficacy by increasing the mean estimate of the area-normalized peak resistance by 0.28 g/mm^2^ (Fig 1H). The addition of GSH had a protective effect suggested by a substantial reduction in the mean area-normalized peak resistance from 2.94 g/mm^2^ to 1.60 g/mm^2^ (−46 %)(Fig 1H). Interestingly, simultaneous addition of the AAPH and GSH showed a synergistic rather than antagonistic effect further decreasing the PASTA-derived estimate to 0.91 g/mm^2^ (31% of the CTR) and supporting the observation that oxidation does not exert a pronounced effect on the lubricating efficacy of the tested lubricant (Fig 1H). Moreover, a fresh sample of the lube demonstrated substantially better lubricating efficacy in comparison with the control sample exposed to air for 120 minutes (1.49 g/mm^2^ vs 2.94 g/mm^2^) with PASTA-derived values resembling those observed with the GSH treatment (Fig 1H). Finally, to test whether the observed changes were associated with other tribological properties the capillary movement and the movement on a clean borosilicate surface placed at a 90° angle were analyzed. The AAPH treatment induced minimal alterations in the capillary movement capacity but demonstrated a pronounced potentiation of the movement on the flat borosilicate surface (Fig 1I-K). Pre-incubation with GSH had a slightly more pronounced effect in the same direction, and simultaneous treatment with both AAPH and GSH exerted a synergistic effect resulting in the greatest change in both assays (Fig 1I-K). The analysis of movement profiles on the vertical flat borosilicate surface demonstrated that all treatments decreased the pinning phenomenon when compared to the control untreated lubricant gel and that the synergistic effect of AAPH and GSH was associated with a slight increment in sliding speed once depinning was achieved (0.41 mm/s [AAPH+GSH] vs 0.36 mm/s [AAPH] and 0.35 mm/s [GSH])(Fig 1K). Considering that the results of the PASTA-based tribometric analysis may have been influenced by the molarity (introduced by the addition of GSH and AAPH) an additional control experiment was conducted in which the effect of the dilution of the lubricant with ddH2O was tested showing that reduced molarity improved lubricating properties (Fig 1L).

## Discussion

The results presented here provide evidence that the modular open-source kitchen scale-based research platform PASTA, originally developed for the analysis of the acoustic startle and sensorimotor gating in rodents [14] and later adapted for the quantification of grip strength [15], can also be used for the assessment of tribological properties. To record forces made by friction between the concave and the convex surface of two complementary PCV tubes, the platform has to be adapted to enable a hollow PVC tube holder to be tightly attached to the bottom of the kitchen scale in a way that makes the tubes freely accessible for the administration of the dynamic and/or static tension. The mastPASTA adapter proposed here was designed to be compatible with the idea of PASTA and open-source bioinstrumentation in general – the adapter is minimalistic and easy to make using only widely available and affordable equipment and its modular design permit further adaptation to meet the requirements of different research challenges.

Considering that a setup presented here demonstrated a satisfactory linear association between the PASTA-estimated peak resistance and the area of the contact surface between the two PVC tubes following the administration of a relatively unreliable dynamic force (i.e. a downward pull of the PVC tube exerted by the researcher), the same approach was used in the proof of concept experiment. Nevertheless, it can be assumed that the administration of a static force can provide more reliable estimates and a deeper insight into different tribological properties of the lubricant being tested. With this in mind, it is emphasized that a more controlled assay can be designed by i) attachment of a standard weight on the bottom of the inner PVC tube (e.g. by using a clip or a hook); or ii) installation of a controlled engine pulley beneath the platform (that can then be interfaced with PASTA for complete automation).

The utility of using mastPASTA and PASTA for the assessment of tribological properties was further tested in the proof of concept study on the effects of redox-related alterations on the lubrication capacity of the commercial water-based lubricant gel. The original hypothesis was that water-based lubricants for personal use lose their lubricating capacity following oxidation. Interestingly, although the addition of AAPH successfully induced oxidation of the lubricant (confirmed by the ORP analysis; Fig 1F) the observed lubricating capacity decrement was minimal and unconvincing (Fig 1H). In contrast, the addition of the antioxidant GSH (associated with the greater antioxidative capacity of the lubricant as shown by ORP) increased the lubricating capacity after exposure to air (Fig 1H). The fact that the simultaneous addition of AAPH and GSH demonstrated a synergistic effect on lubricating capacity (Fig 1H) regardless of the fact that it didn’t affect ORP (Fig 1F) provides additional evidence to reject the original hypothesis – in contrast to what was expected the oxidation does not seem to affect the lubricating properties and the protective effect of GSH does not seem to be dependent on its antioxidant properties. One possible explanation of the observed protective effects of GSH may be the altered pH of the lubricant as it is a well-known fact that acidity can have a profound effect on lubrication properties[33,34]. Nevertheless, the fact that the addition of both AAPH and GSH resulted in a siminar modest change in acidity speaks against the pH acting as a mediator of the phenomenon observed here. Another possible explanation unrelated to redox changes may be that addition of AAPH, GSH, and in particular, the combination of AAPH and GSH altered the lubrication properties by simply increasing the osmolality of the lubricant. This explanation is in line with other biotribometric experiments as the lubricant with both AAPH and GSH shows the shortest capillary movement (Fig 1I) and the longest distance on the flat borosilicate surface at the 90° angle (Fig 1J). Nevertheless, examination of lubrication efficacy of the ddH2O-diluted lubricant (Fig 1L) shows that increased osmolarity of the tested lubricant would most likely be associated with decreased and not increased lubricating capacity. Consequently, while increased molarity may provide an explanation for the observed alterations in capillary movement and movement on the flat glass surface it does not explain the improved lubricating properties of the GSH-treated lubricant in the lubrication efficacy PASTA-based biotribometric assay. The latter observation provides an additional invaluable insight because many commercially-available water-based vaginal lubricants are characterized by high osmolarity with the idea that hyperosmolar products will attract water and facilitate lubrication through increased volume. Unfortunately, it has been shown that hyperosmolarity is associated with mucosal irritation [35] and damage to the vaginal epithelial barrier[36], the effect not observed with iso and hypo-osmolar vaginal lubricants. As increased osmolarity seems to unfavorably affect the lubricating properties of the water-based lubricant gel tested here the question arises whether the increased osmolarity of lubricating gels is associated with better lubrication *in vivo.* If this is not the case (as osmolarity attracts water, but also negatively affects the intrinsic lubricating potential) hyperosmolar lubricants should be reevaluated as they may convey no additional benefits while increasing epithelial damage[35,36] and the risk of sexually transmitted diseases[37].

## Limitations

The research presented here was designed as a proof of concept for the use of PASTA for testing biotribometric properties. For this reason, the experiment on the effects of redox-related alterations on the lubrication capacity of the commercial water-based lubricant gel is relatively simple and associated with several limitations. First, only a single lubricant gel was tested. Consequently, although it can be assumed that most of the observed effects are applicable to different lubricants, there is a possibility that the randomly selected lubricant does not reliably represent the whole population of vaginal lubricants (due to variable composition, pH, osmolarity, etc.). Second, although several measures were introduced to reduce the risk of bias (e.g. evaporation monitoring, ORP, and pH assessment, etc.) additional control experiments could have been conducted. For example, to reliably assess the effects associated with pH, an additional experiment could have been introduced with all tested samples adjusted to the original pH value of the standard lubricant. As there was indirect evidence that pH is not the mediator of the observed effects the introduction of this experiment would dramatically increase the required time and resources and contribute nothing to the general goal of the study – to demonstrate the use of PASTA for biotribometric analysis. Additionally, the same experiment could have been repeated with different concentrations of AAPH and GSH, and different dilutions of the lubricant (to simulate the attraction of water expected to happen *in vivo*). Although there is no doubt that the aforementioned experiments would provide exciting new data they will be conducted on another occasion.

## Conclusion

To conclude, the PASTA platform with the mastPASTA adapter provides an easy, affordable, and open-source solution for reliable quantitative measurements in simple biotribometric assays. In contrast to the original hypothesis, the oxidation does not seem to dramatically affect the lubrication properties of the commercial water-based lubricant gel. Nevertheless, the addition of the antioxidant GSH seems to increase the lubricating properties of the gel and/or its resilience to the effects of air exposure by redox-unrelated mechanisms that remain unresolved. Considering the importance of oxidative stress in the etiopathogenesis of vulvovaginal dryness, the addition of GSH (and other antioxidants) should be explored as a possible way to enhance the durability and efficacy of vaginal lubricants and increase their health benefits.

## Conflict of interest

None.

## Funding

None.

## Notes

### Competing Interest Statement

The authors have declared no competing interest.

### Summary of Updates

Updated figure 1

